# mettannotator: a comprehensive and scalable Nextflow annotation pipeline for prokaryotic assemblies

**DOI:** 10.1101/2024.07.11.603040

**Authors:** Tatiana A. Gurbich, Martin Beracochea, Nishadi H. De Silva, Robert D. Finn

## Abstract

**Summary:** In recent years there has been a surge in prokaryotic genome assemblies, coming from both isolated organisms and environmental samples. These assemblies often include novel species that are poorly represented in reference databases creating a need for a tool that can annotate both well-described and novel taxa, and can run at scale. Here, we present *mettannotator* - a comprehensive Nextflow pipeline for prokaryotic genome annotation that identifies coding and non-coding regions, predicts protein functions, including antimicrobial resistance, and delineates gene clusters. The pipeline summarises the results of these tools in a GFF (General Feature Format) file that can be easily utilised in downstream analysis or visualised using common genome browsers. *Mettannotator* has been tested on both novel and known taxa, and used across Ensembl Bacteria and MGnify. Here, we show how it works on 29 prokaryotic phyla, both isolate and metagenome-assembled genomes, and present metrics on its performance in comparison to other tools.

**Availability and implementation:** The pipeline is written in Nextflow and Python and published under an open source Apache 2.0 licence. Instructions and source code can be accessed at https://github.com/EBI-Metagenomics/mettannotator. The pipeline is also available on WorkflowHub: https://workflowhub.eu/workflows/1069.

## 1 Introduction

The rapid rise of affordable high-throughput sequencing has resulted in a deluge of microbial genomes from isolates, single-cell sequencing and metagenomes (Hunt et al. 2024). While there are numerous genome annotation tools available, it is not always easy to install them, run them at scale, or merge their outputs into a single useful format - a major drawback for teams without bioinformatics expertise. Furthermore, while many pipelines hone in on the annotation of individual proteins, there is lesser focus on the identification of larger gene clusters which provide contextual information, the systematic annotation of antimicrobial resistance genes, or collating the results of different tools, to provide deep insights into the functional role of a microbe within its community (Jung et al. 2024). Annotation of novel genomes presents an additional challenge as these genomes are less likely to be represented in reference databases which may hinder the computational transfer of functional information.

To address these issues, we have developed a comprehensive pipeline - *mettannotator* - that combines existing tools and custom scripts to perform both structural (demarcating genomic elements) and functional (assigning functions to genomic elements) annotation of prokaryotic genomes. By employing a diverse set of annotation tools, the pipeline is able to handle genomes even without a species-level taxonomic label. In addition to using several reference databases, *mettannotator* builds upon established annotation frameworks used in UniProt (The UniProt Consortium et al. 2023) to assign function to as many unannotated proteins as possible. It also predicts larger genomic regions such as biosynthetic gene clusters, anti-phage defence systems and putative polysaccharide utilisation loci, and consolidates all annotations into a final GFF file. Implemented in Nextflow and nf-core (Ewels et al. 2020; Di Tommaso et al. 2017) and following the best practices established by the nf-core community, *mettannotator* is fully containerised, making it portable and easily deployable, scales well, is versioned (ensuring annotation provenance), and is discoverable. The pipeline can be run on isolate and metagenome-assembled genomes (MAGs), both bacterial and archaeal, and is now the unified annotation pipeline used by several microbial resources, namely Ensembl Bacteria and MGnify (Harrison et al. 2024; Gurbich et al. 2023). We present the results of our pipeline on 200 genomes and compare it to other prokaryotic annotation packages.

## 2 Pipeline description

### 2.1 Installation and dependencies

Nextlow and Singularity/Apptainer or Docker are the only prerequisites for running *mettannotator*. No tool installation is required. Databases are downloaded automatically during the first execution of the pipeline unless provided by the user. The total size of the reference databases is 180 GB. Although it is possible to run the workflow on a personal computer, it is recommended to execute it on a compute cluster for large scale annotation efforts. A minimum of 50 Gb of RAM and 16 CPUs are required to run *mettannotator*.

### 2.2 Input files

The pipeline takes a single comma-separated text file as input which can include one or many genomes to be analysed. For each genome, it is required to provide:

- a prefix, which will be used to name the result files and in the locus tags for features in the GFF file
- a path to the genome assembly in standard FASTA format (ideally oriented to start at dnaA or repA)
- a valid NCBI TaxId for the lowest known taxonomic level

### 2.3 Annotation workflow

The workflow schema is shown in Figure 1A. The *mettannotator* pipeline offers the user a choice between two well established prokaryotic gene prediction tools - Prokka (Seemann 2014), the default choice, and Bakta (Schwengers et al. 2021). Since Bakta is only intended for bacterial genome annotation, *mettannotator* detects the domain automatically and always uses Prokka to predict genes in archaeal genomes.

**Figure 1:**
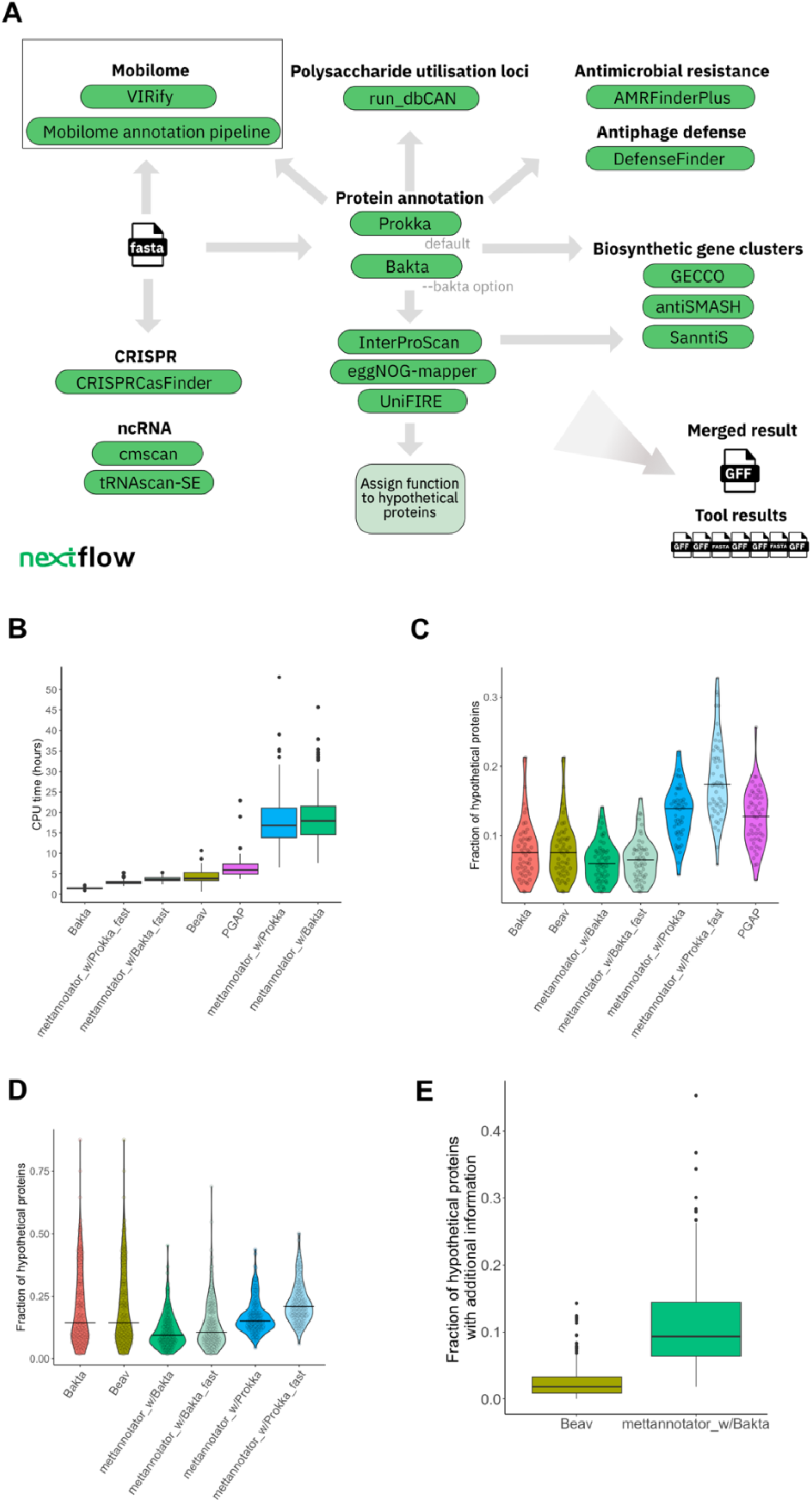
**A**) The *mettannotator* workflow schema. The pipeline outputs the results of the individual tools as well as a merged GFF file. The mobilome annotation is not currently integrated into the Nextflow pipeline but can be executed separately. The results can be added to the *mettannotator* GFF output file using a provided post-processing script. **B**) CPU time per genome. *Mettannotator* (with Bakta and with Prokka as the gene caller, with and without the --fast flag) was run on 200 genomes , Bakta and Beav on 194 bacterial genomes, and PGAP on 51 genomes with known species-level taxonomy confirmed by PGAP. *Mettannotator* was faster than both PGAP and Beav when executed in the fast mode. When executed without the fast flag, *mettannotator* took longer than other tools to complete. **C**) Fraction of hypothetical proteins out of all detected CDS, calculated for the 51 genomes that were annotated by PGAP demonstrating tool performance on genomes with known and confirmed species-level taxonomy. Horizontal bars indicate the median value. *Mettannotator* with Bakta as the gene caller has the lowest fraction of hypothetical proteins compared to other tools. **D**) Fraction of hypothetical proteins out of all detected CDS. 194 bacterial genomes were used as inputs. When genomes from novel taxa are added to the analysis, the difference between *mettannotator* and other tools is more pronounced. **E**) Fraction of proteins that are labelled as “hypothetical” but have additional information available in the GFF file as reported by Beav and *mettannotator* (run in normal mode with Bakta as the gene caller). Both tools were run on 194 bacterial genomes. Additional information can help the user identify the function of a protein in cases where it was not possible to assign it computationally.

Functional information provided by the gene caller tool is further supplemented by running predicted proteins through InterProScan (Jones et al. 2014), eggNOG-mapper (Cantalapiedra et al. 2021), and UniFIRE, the UniProt Functional annotation Inference Rule Engine (‘UniFIRE’ 2024) . Running UniFIRE allows *mettannotator* to deploy the automated function prediction system used by UniProt based on three sets of UniProt rules: curated UniRule and UniRule-PIRSR and automatic, rule-mining based ARBA (MacDougall et al. 2020; Chen et al. 2019; Saidi et al. 2017).

After this, any remaining proteins that are labelled as *hypothetical* are further processed to select probable function from the outputs of InterProScan, UniFIRE and eggNOG-mapper. The function assignment logic and prioritisation of data sources are shown in Supplementary Figure 1. When functional assignment is possible, the “product” field in the final GFF file produced by *mettannotator* is overwritten to reflect the detected function.

Furthermore, *mettannotator* predicts antimicrobial resistance genes, virulence factors, biocide, heat, acid and metal resistance genes by running AMRFinderPlus (Feldgarden et al. 2021) and known antiviral defence systems using DefenseFinder (Tesson et al. 2022). To provide the user with the widest range possible for potential biosynthetic gene clusters, the pipeline runs antiSMASH (Medema et al. 2011), GECCO (Carroll et al. 2021), and SanntiS (Sanchez et al. 2023) and reports the outputs of each tool. Furthermore, CAZyme gene clusters (which if experimentally validated can be categorised as polysaccharide utilisation loci (PULs)) and their putative substrates are annotated using run_dbcan, the standalone version of dbCAN3 (Zheng et al. 2023), and CRISPR arrays are predicted using CRISPRCasFinder (Couvin et al. 2018). We use tRNAscan-SE (Chan and Lowe 2019) for tRNA annotation and cmscan (Nawrocki and Eddy 2013) against the Rfam database (Kalvari et al. 2021) to identify and annotate rRNA and other non-coding RNA. Versions of all tools and databases are listed in the README file in the GitHub repository, and reported by the pipeline in the MultiQC (P. Ewels et al. 2016) summary report.

*Mettannotator* also provides a convenient option to generate quick, draft annotations by using the ‘--fast’ flag. These are less in-depth, but take a fraction of the time as the pipeline skips InterProScan, UniFIRE and SanntiS. Functions of fewer proteins are resolved when the --fast flag is used as eggNOG-mapper becomes the only additional source of information for the “product” field.

Nextflow allows for seamless parallel annotation of multiple genomes, monitoring and restarting the pipeline from any points of failure. At the end, the pipeline parses the results of each step and consolidates them into a final GFF file per genome.

### 2.4 Output files

The *mettannotator* pipeline produces a final valid GFF file with results from the above tools merged. The ninth column of the file contains carefully chosen key-value pairs to report the salient conclusions from each tool. Notably, we include KEGG orthology identifiers (Kanehisa et al. 2017), gene ontology (GO) terms (Thomas 2017) , and Chemical Entities of Biological Interest (ChEBI) identifiers (Degtyarenko et al. 2007) parsed from eggNOG-mapper and UniFIRE outputs.

The annotations are visualised using the PyCirclize package (‘pyCirclize’ 2022) to produce a Circos plot (Krzywinski et al. 2009) for each genome with no more than 50 contigs. Additionally, the pipeline output includes key files generated by each tool, organised in a structured directory system for each genome, facilitating deeper analysis. The source code and documentation for the pipeline is available via GitHub.

### 2.5 Mobilome annotation

The GFF file produced by *mettannotator* can be used as input for our mobilome annotation pipeline (‘Mobilome Annotation Pipeline’ 2024) - a separate, yet compatible workflow that, in concert with VIRify (Rangel-Pineros et al. 2023), a viral sequence detection and annotation tool, annotates plasmids, phages and integrative elements. We provide a post-processing Python script that can be executed separately to incorporate the results of the mobilome annotation pipeline back into the GFF file produced by *mettannotator*.

## 3 Performance evaluation

### 3.1 Test dataset and tools

To assess *mettannotator*’s performance in comparison to existing tools, we put together a test dataset that included 200 genomes: *E*.*coli* O26:H11 str. 11368 genome from GenBank (GCA_000091005.1) (Ogura et al. 2009) and 199 genomes from MGnify (Gurbich et al. 2023), both MAGs and isolates, bacteria and archaea, from 6 different biomes: human gut, human oral, cow rumen, fish gut, marine, and chicken gut. The genomes were chosen to cover a wide range of taxa, including uncharacterised ones, and different levels of genome completeness, contamination and contiguity (Supplementary Table 1).

We annotated this dataset using *mettannotator* v.1.2 with Prokka and with Bakta as the gene callers (executed with and without the --fast flag), Bakta v1.9.3 (Schwengers et al. 2021) , PGAP v.2024-04-27.build7426 (Haft et al. 2024; Li et al. 2021), and Beav v1.3.0 (Jung et al. 2024). Bakta and PGAP were chosen for comparison as these are two of the more widely used annotation tools and Beav was added as this workflow is similar to *mettannotator*. The tool execution parameters are described in Supplementary Methods. Only annotation types generated in at least one other workflow in addition to *mettannotator* were compared.

We introduced several constraints when executing third-party pipelines. The archaeal genomes were excluded when using Bakta and Beav as these tools are only intended for bacterial genomes. Beav was executed with --skip-tiger flag, skipping integrative conjugative element (ICE) analysis, since we were unable to find the expected reference database or how to generate it. When performing annotation with Bakta and Beav, we skipped CRISPR annotation for two genomes due to Bakta errors that lead to execution failure, however, these genomes were retained for the comparison. Finally, since PGAP requires that at least the genus of the genome being annotated is known, we were only able to annotate 51 out of 200 genomes with PGAP. The taxonomies of the remaining genomes were either labelled as inconclusive or the genome was marked as contaminated by PGAP.

### 3.2 Performance results

First, we compared per genome CPU time between all tools (Figure 1B). Each tool was run on all genomes that it was able to process: a full set of 200 genomes for *mettannotator*, bacterial genomes for Beav and Bakta (194 genomes), and 51 genomes with confirmed species-level taxonomy for PGAP.

When executed with the --fast flag (skipping InterProScan, UniFIRE and SanntiS), the average *mettannotator* CPU time was 3.77 hours with Bakta as the gene caller and 2.96 hours with Prokka, which is faster than any tool except Bakta executed on its own. *Mettannotator* consumed more resources per genome compared to the other tools when the full version was executed, with an average CPU time of 18.71 hours when using Bakta and 18.04 hours when using Prokka. The execution of InterProScan and UniFIRE were the most resource-intensive steps (Supplementary Figure 2). However, these tools provide extensive functional information and are used by *mettannotator* to identify functions of unannotated proteins.

To measure the wall-clock time and provide guidance to end users, we benchmarked *mettannotator* using its two gene-calling tools, Prokka and Bakta, in both normal and fast modes. We annotated 8 MAGs from the MGnify Genomes collection. Additionally, we compared the results with Bakta v1.9.3 using its default settings. The benchmark was performed on a virtual machine equipped with 32 vCPUs (Intel Xeon Cascade Lake 2.5GHz) and 128GB of RAM.

Bakta completed the annotations with an average wall-clock time of 16 minutes ± 6 minutes per genome. In comparison, *mettannotator*, when utilising Prokka in normal mode, averaged 64 minutes ± 14 minutes. However, when *mettannotator* used Prokka in fast mode, the average wall-clock time decreased to 12 minutes ± 5 minutes. Using *mettannotator* with Bakta in normal mode resulted in the longest average wallclock time of 97 minutes ± 14 minutes. In fast mode, *mettannotator* with Bakta averaged 41 minutes ± 8 minutes.

To compare annotation results between the tools, we evaluated the fraction of unannotated proteins per genome. We refer to these proteins as “hypothetical” in line with their product label. We classified any CDS with either “hypothetical protein” or “uncharacterized protein” listed as the product in the ninth column of the final GFF file as hypothetical. While some of these proteins might have additional annotations available, the tool generating the annotation could not assign a product due to either a lack of a high quality database hit or the specifics of the annotation algorithm.

We performed initial comparison using only the 51 genomes that we executed PGAP on (Figure 1C). These genomes have known species-level taxonomy and high level of completeness, with an average of 95.65% as calculated by CheckM (Parks et al. 2015).

All tools produced few hypothetical proteins in this comparison, with the median fraction of hypothetical proteins in a genome below 20%. Bakta, Beav and *mettannotator* with Bakta as the gene caller had fewer hypothetical proteins than PGAP. *Mettannotator* had the lowest fraction of hypothetical proteins out of all tools when using Bakta as the gene caller (median = 5.9%). Bakta and Beav show the same result in this comparison since Beav does not override the product name assigned by Bakta, instead adding insights from other tools into the “Note” field of the GFF.

We then ran all of the tools except for PGAP on 194 bacterial genomes. In this comparison, which included genomes with unknown taxonomy, the difference between the fraction of hypothetical proteins in *mettannotator*’s results and the results of the other tools was more pronounced (Figure 1D). When run with Bakta as the gene caller, *mettannotator* results had fewer hypothetical proteins than any of the other tools (median = 9.3%). The difference between *mettannotator* and other tools is even greater when run on a subset of genomes with no assigned species name or with poorly known taxonomy (Supplementary Figure 3), showing that *mettannotator* can be particularly useful when annotating novel genomes.

To ensure that the sources of protein function are reliable, *mettannotator* uses well-established databases and approaches, including UniFIRE and InterPro that are utilised by UniProt. The majority of hypothetical proteins relabeled by *mettannotator* inherit function from InterPro (Supplementary Figure 4).

Lastly, we looked into additional functional information that is reported by Beav and *mettannotator* but is not included in the product description. In the Beav annotation format, additional functional information is included in the “Note” field. In the *mettannotator* GFF file there can also be cases where the product field states that a protein is hypothetical, however, additional tools provide an annotation. To account for this, we looked at the fraction of hypothetical proteins that have additional information from other tools available (Figure 1E). *Mettannotator* has a higher fraction of such proteins, indicating that although in some cases we are not able to assign a function, additional information provided, such as, for example, an antimicrobial or antiviral defence annotation, can lead the user towards a possible role of the protein.

## 4 Conclusion

*Mettannotator* offers a comprehensive annotation of prokaryotic genomes. It is able to assign a function to more proteins than existing workflows, however, does so at a slight cost of compute time and resources. *Mettannotator* can be particularly useful when processing novel genomes, including batch-processing large numbers of genomes, as long as their taxonomic domain is known. Several features of *mettannotator* are not usually found in similar pipelines, for instance, the chemical reaction information provided through ChEBI identifiers and functional annotations by UniFIRE.

*Mettannotator* has already been taken up by several public resources. An earlier version of this pipeline was used to re-annotate all prokaryotic genomes in Ensembl Bacteria (∼32,000) in 2023 as well as the genomes in the MGnify genome catalogues.

## Acknowledgements

The authors would like to thank Carlos Voogdt and Mariia Beliaeva from EMBL Heidelberg, Indra Roux and Alexandre Almeida from the University of Cambridge, Olivia Casanueva, Ekaterina Sakharova, Alejandra Escobar, and Evangelos Karatzas at EBI Hinxton for various invaluable input towards the construction of this pipeline.

## Data availability

The pipeline is freely available at https://github.com/EBI-Metagenomics/mettannotator

