## Supplementary Material for "mettannotator: a comprehensive and scalable Nextflow annotation pipeline for prokaryotic assemblies"

### Supplementary methods

#### Test dataset

Completeness and contamination of the 200 genomes included in the test dataset were calculated using CheckM v1.1.3 (Parks et al. 2015). Taxonomy was determined using GTDB-Tk v.2.4.0 with reference database release 220 (Chaumeil et al. 2020). TaxId for the *mettannotator* input file was determined using CAT v.5.3 (von Meijenfeldt et al. 2019). *E.coli* (GCA_000091005.1) taxonomy was manually replaced as CAT only classified this genome down to the superkingdom level.

The dataset includes 159 MAGs and 41 isolate genomes. The average genome completeness is 89.2% and maximum contamination is 5.13%. The number of contigs ranges from 1 to 889 and the genome length ranges from 583.5 Kb to 10.6 Mb. Twenty-nine prokaryotic phyla are represented in the dataset.

#### Tool execution

*mettannotator*

*Mettannotator* was run in 4 modes - with Prokka and with Bakta as the gene callers and with and without the --fast flag. To compute CPU time, we executed *mettannotator* one genome at a time and used the CPU time reported by Nextflow.

PGAP

PGAP requires that the user provides either the genus or the species name as the organism name when running the annotation workflow. To determine taxonomies that would pass PGAP's checks, we first executed PGAP on each test genome with the --taxcheck-only flag enabled. Out of 200 genomes, PGAP found a "best match" for 137 genomes. We reran PGAP on these genomes with the --taxcheck-only flag enabled providing PGAP's highest species-level taxonomy match from the results of the previous run as the organism name. Taxonomy of 42 genomes was confirmed by PGAP with high confidence, further 9 genomes were confirmed with low confidence, and two MAGs, MGYG000299175 and MGYG000299211, were marked as contaminated by PGAP but reported as having high confidence. We included all of the genomes with confirmed taxonomy, 51 in total, in the annotation input dataset. We were not able to improve PGAP's taxonomy confirmation result when we only provided the genus name rather than the species name for any of the genomes in our dataset. Consequently, all genomes that were retained for further analysis were given species-level organism names.

The dataset annotated by PGAP comprises 25 MAGs and 26 isolate genomes. It includes 22 genomes from the human gut, 8 from the human oral cavity, 7 from marine environments, 7 from the fish gut, 6 from the chicken gut, and 1 from the cow rumen. The average genome completeness is 95.65% (Supplementary Table 1).

Genomes were annotated using 16 CPUs and 50 Gb of RAM and with --no-internet, --no-self-update and --ignore-all-errors flags enabled. The Docker container was downloaded prior to execution and no tax check was performed during the annotation process.

Bakta

Bakta was run on all bacterial genomes in the test dataset (194 in total) using 8 CPUs and 25 Gb of RAM per genome. Two genomes (MGYG000298952 and MGYG000298476) were annotated with the --skip-crispr flag enabled due to this portion of the workflow failing.

Beav

Beav was run on all bacterial genomes with 8 CPUs and 32 Gb of RAM per genome and the --skip_tiger flag. We built a Singularity container that included the databases Beav needs to execute the workflow. Same as with Bakta, 2 genomes (MGYG000298952 and MGYG000298476) were annotated with the --skip-crispr flag enabled due to this portion of the workflow failing.

When performing CPU time calculations, CPU times for Beav, Bakta and PGAP were obtained from Slurm reports.

**Comparing additional information available for hypothetical proteins**

In Figure 1E we compared the number of hypothetical proteins with additional information between Beav and *mettannotator* (run in normal mode with Bakta as the gene caller). We ran both tools on all bacterial genomes in the test dataset (194 genomes). We considered proteins that have “hypothetical protein” or “uncharacterized protein” as product to be hypothetical. To identify the number of hypothetical proteins that have additional information in the output of Beav, we counted hypothetical proteins that have a “Note” field in the ninth column of the GFF file. To identify the number of hypothetical proteins with additional information in the output of *mettannotator*, we counted the number of hypothetical proteins that have at least one of the following fields in the ninth column of the GFF file: “Ontology_term”, “amrfinderplus_gene_symbol”, “amrfinderplus_scope”, “amrfinderplus_sequence_name”, “cog”, “dbcan_prot_family”, “dbcan_prot_type”, “drug_class”, “drug_subclass”, “eggNOG”, “interpro”, “kegg”, “pfam”, “substrate_dbcan-pul”, “substrate_dbcan-sub”, “uf_chebi”, “uf_gene_name”, “uf_gene_name_synonym”, “uf_keyword”, “uf_ontology_term”, “uf_pirsr_cofactor”, “uf_prot_alt_ecnumber”, “uf_prot_alt_fullname”, “uf_prot_alt_shortname”, “uf_prot_rec_ecnumber”, “uf_prot_rec_fullname”, “uf_prot_rec_shortname”, “note”.

**Supplementary Figure 1:**

Algorithm to assign functions to proteins labelled as hypothetical by the gene caller. UniFIRE, InterProScan and eggNOG-mapper results are used as sources of functional information.

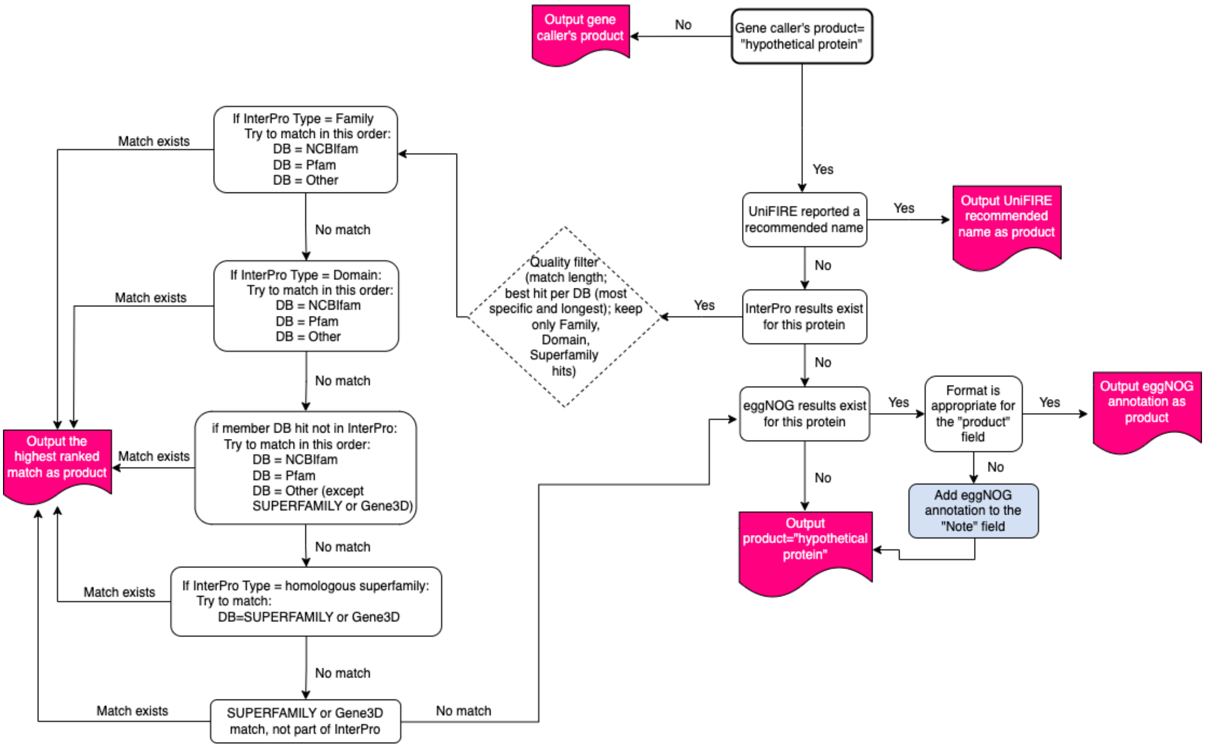

**Supplementary Figure 2:**

Amount of compute time per process when annotating 194 bacterial genomes using *mettannotator* with the --bakta flag enabled.

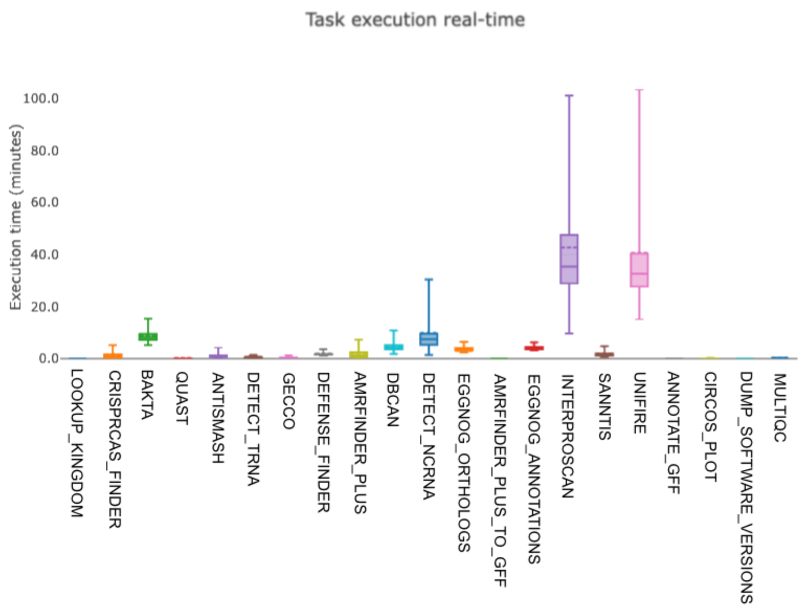


**Supplementary Figure 3:**

Fraction of hypothetical proteins in genomes with poorly known taxonomy. Tools were run on 61 genomes (a subset of the 200 genomes used for benchmarking) that either did not have a species name according to GTDB taxonomy (Supplementary Table 1) or did not use a typical binomial nomenclature indicating the genome likely has not been isolated (for example, "*CAAFZY01 sp900767645*" or "*Phil1 sp004558525*"). *Mettannotator* with Bakta as the gene caller run in normal mode had the least median fraction of hypothetical proteins per genome.

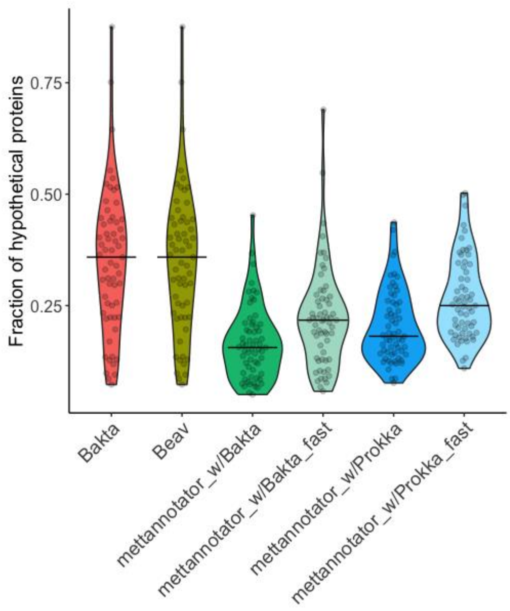

**Supplementary Figure 4:**

The source of product description for CDS in each genome when replacing a protein labelled as "hypothetical" by the gene caller (here, Bakta was used as the gene caller for bacterial genomes). InterPro is used more often to relabel the product than UniFIRE or eggNOG. When AntiFam is the source of product description, the product is likely a spurious ORF.

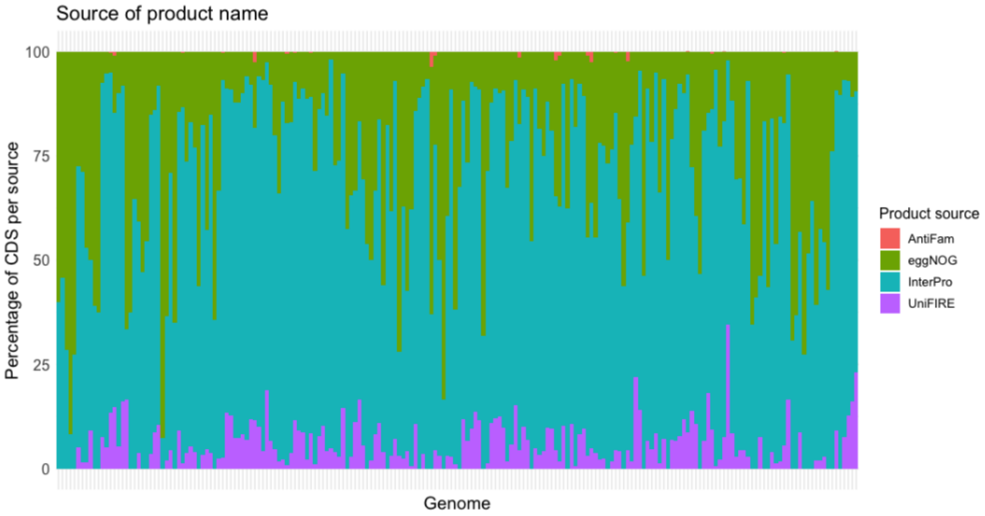
